# Transmission strategy modulates parasite biogeography in an island-colonising bird

**DOI:** 10.64898/2026.01.30.702753

**Authors:** Sarah Nichols, Andrea Estandía, Fiona Robertson, Bruce C. Robertson, Beth Okamura, Sonya M. Clegg

## Abstract

Parasites occur in every ecosystem, although their dispersal is often constrained by the availability of hosts or vectors. Here, we explore how variation in parasite life history traits, particularly transmission strategy, may influence their distributions. Specifically, we test whether a variety of parasites adhere to the rules of island biogeography, and whether their distributions vary with transmission strategy. We utilised broad-spectrum parasite detection from existing Whole Genome Sequence (WGS) data to characterise parasites with varying transmission strategies from the blood of a passerine bird, the silvereye (*Zosterops lateralis*), sampled across 25 islands in the South Pacific and from five of the states in mainland Australia. Overall, parasite richness was higher on mainland Australia compared to islands and decreased with distance of islands from the Australian continent. However, these patterns were dependent on transmission strategy. For parasites transmitted by flying insect vectors, richness decreased on islands compared to the mainland. However, increasing isolation from the mainland among islands had little further impact. On the other hand, richness of directly transmitted parasites and those requiring another intermediate host declined sharply with increasing distance from the mainland. While islands may act as an initial barrier to colonisation for parasites relying on flying insect vectors, their highly dispersive vectors may subsequently reduce the impact of increasing isolation distance on richness. Our work underscores the importance of considering parasite life-histories and their transmission strategies for understanding the processes that shape parasite communities on islands.

## Introduction

Parasitic taxa can account for up to 40% of animal biodiversity in an ecosystem (1–3) and can thus have profound and varied impacts on ecosystem dynamics (4). They may influence inter- and intra-specific interactions, diversification rates and ultimately global patterns of biodiversity (5–8). Therefore, characterising the biogeographic patterns of parasites is important. However, parasites rely on hosts for essential resources (9, 10), their distributions are subsequently dictated by host availability and movement. Parasite dispersal can be further complicated by their own life histories, including transmission strategy. How such life history features influence the biogeographic distributions of parasites is unclear and exacerbated by our generally poor knowledge of parasite diversity (11, 12) and difficulty in defining the geographic boundaries of their distributions (13). To address these limitations, application of molecular methods to provide broad-spectrum characterisation of parasite communities in an island biogeography framework offers a powerful means for advancing our understanding of how life history traits shape parasite distributions.

The rules of island biogeography are based on the premise that a mainland (e.g. a continental region or large land mass) acts as a source for islands. A subset of species is expected to colonise and persist on islands, resulting in lower equilibrium species richness on islands, with the lowest richness on small, remote islands, following species-area and distance-decay relationships (14). While the framework was developed for free-living organisms, parasite distributions are assumed to behave similarly because parasites are closely associated with their hosts; therefore, island features will similarly constrain colonisation and extinction processes (15). Studies have shown the expected pattern of lower parasite diversity on islands for various ecto- and endoparasites in birds (16), amphibians (17), fish (18) and mammals (19). However, conflicting patterns have also been reported in a similarly broad range of parasites and hosts, including birds (20), amphibians (21), and insects (22). Even within one group of well-studied parasites, avian haemosporidians, deviations from island biogeography expectations are frequently reported. For instance, variation in distance-decay relationships has been observed among haemosporidian genera in South Pacific island white-eyes (Zosteropidae) (23) and for haemosporidian lineage richness patterns in birds of the Gulf of Guinea (24). These deviations from island biogeography expectations are often attributed to differences in parasite life histories, such as life cycle complexity (10, 25), or level of host specialisation (23).

Transmission strategies are a potentially important life history trait to consider when aiming to understand patterns of parasite distributions. Parasites with simple life cycles utilise direct transmission strategies that include host contact, faecal-oral routes and environmental contamination (26). Many directly transmitted parasites can produce encysted stages, with a resistant outer layer that provides protection from adverse environmental conditions, such as temperature extremes and desiccation (27, 28). Encysted stages can persist in the environment when suitable hosts are unavailable, increasing the chance of establishment (29, 30).

These dormant stages may also be dispersed via zoochory when attached to mobile animals (31). Directly transmitted parasites spend the majority of their replicating life cycle within a host (27) and are likely to co-disperse with them, thus their patterns of dispersal may reflect that of their hosts (32). As birds show strong adherence to island biogeography expectations (33), it follows that directly transmitted parasites of birds will likewise show such patterns.

Parasites with complex life cycles are characterised by a range of transmission strategies. Some are transmitted via bites from ticks or flying insect vectors, others are transmitted trophically, via consumption of additional infected hosts. The ability of such parasites to colonise islands may vary. For example, infected ticks may co-disperse with birds across extensive distances (34–36). Colonisation of islands by parasites disseminated by flying insect vectors may depend on environmental factors such as offshore wind (37). Establishment of parasites with complex life cycles may also be more challenged by environmental variation than is the case for directly transmitted parasites (38). Reliance on multiple hosts may make persistence more precarious as outcomes will depend on local availability of appropriate additional hosts or vectors (39–41).

To date, most studies on the patterns of island biogeography in parasites have examined single parasite taxa (42–47). This limits our understanding of how parasite life history traits, such as transmission mode, may impact biogeographic patterns. However, characterising a wider diversity of parasites is challenging, particularly because specific host-parasite interactions are not well documented (48, 49). This makes it difficult to select parasite taxa to target. In addition, identification of a broad range of taxa requires expertise in taxonomy, varied methodological approaches, and can incur a high financial cost (4, 11, 50). The ability of high-throughput sequencing technologies to generate Whole Genome Sequence (WGS) data at scale (51) can ameliorate these challenges and offers a new avenue for characterising parasites. A relatively neglected feature of WGS data is that they can also capture information about inadvertently sequenced pathogens and parasites infecting the host (52, 53). Therefore, in addition to addressing evolutionary questions about free-living organisms, WGS data present an opportunity to simultaneously investigate endoparasites of the host. Utilising WGS data in this way enables the identification of a broad range of affiliated parasites with varied life-histories (53).

Here we use WGS data of individual silvereyes, *Zosterops lateralis*, a passerine species that has colonised many islands throughout the South Pacific (54–56), to identify inadvertently sequenced parasites. Our aim is to characterise the distribution and diversity of such parasites in island and mainland silvereye populations to illustrate patterns of island biogeography with respect to richness and nestedness of parasite communities across host populations and to explore how transmission strategy may modulate these patterns. We expect that distributions of parasites utilising direct transmission will more closely reflect island biogeography patterns seen in free-living organisms than those with more complex transmission strategies.

## Materials and Methods

### Study system

The silvereye is a small passerine bird distributed in Australia and islands of the south Pacific (Fig. 1). Their distribution spans a latitudinal range of -14°to -47°. Island silvereye populations vary in age from hundreds to hundreds of thousands of years (56, 57). They occur on islands ranging in size and geographic isolation, from the large Gondwanan islands of Grand Terre, New Caledonia and North and South Islands of New Zealand, to small volcanic islands, such as those of Vanuatu, and coral cays, such as Heron Island on the Great Barrier Reef.

**Fig. 1.**
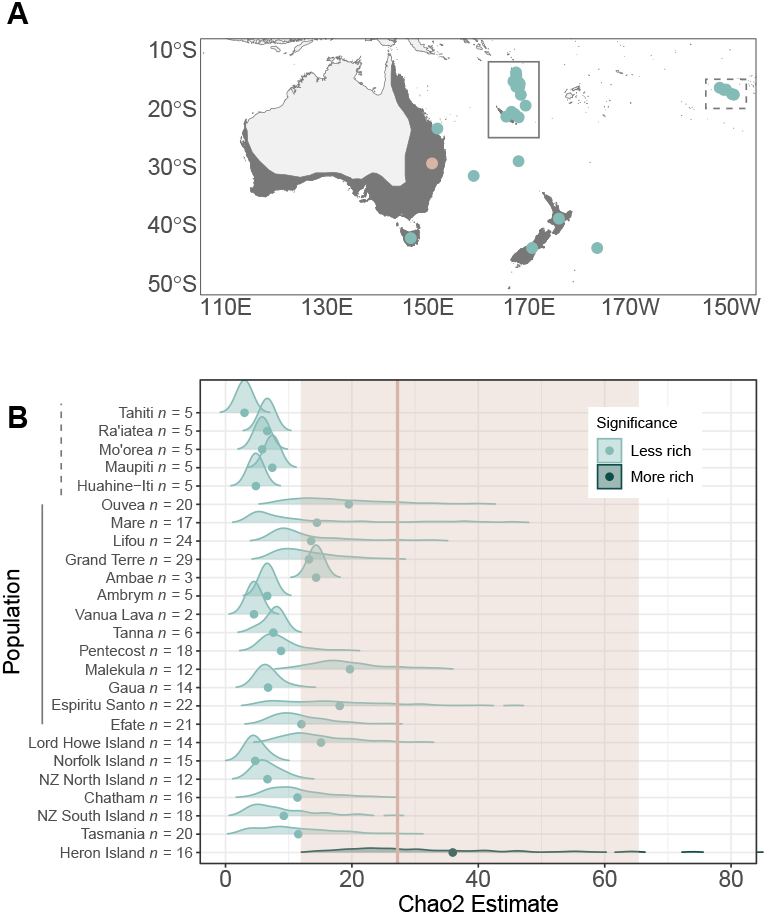
A) Map of sampled mainland (beige) and island (turquoise) populations with dark grey shading indicating the sampled silvereye range. B) Chao2 richness estimates using random-subsampling with replacement to n = 5 with 1,000 bootstraps; the colours of each distribution indicate whether Chao2 richness was significantly more rich (darker) or less rich (lighter) from the mean Chao2 estimate on the mainland, indicated by the beige vertical line and shaded 95% Confidence Intervals (CI).

### Sample collection, DNA extraction and sequencing

The blood samples used in this analysis were collected between 1996 and 2019 from 381 individual silvereyes spanning 25 South Pacific islands and locations on the Australian continent (Queensland, New South Wales, Australia Capital Territory, South Australia and Victoria). Wild birds were caught using mist nets or traps. Blood was taken using a sterile needle to prick the brachial vein. Blood was collected in a glass capillary tube and placed in a 1.5 ml microfuge tube. The microfuge tubes contained Queen’s Lysis Buffer modified from Seutin *et al*. (58) with the final concentrations 0.01 M Tris, 0.01 M NaCl, 0.01 M EDTA and 0.03 M n-lauroylsarcosine (10% w/v), and a pH of 7.5. Samples were kept at ambient temperature until they could be refrigerated and then frozen at -20°C.

DNA extractions were carried out using a modified phenol-chloroform protocol (59). Approximately 100 µl of blood preserved in Queen’s Lysis Buffer was incubated overnight with 250 µl DIGSOL extraction buffer (0.02 M EDTA, 0.05 M Tris-HCl [pH 8.0], 0.4 M NaCl, 0.5% sodium dodecyl sulphate [SDS]) and 10 µl Proteinase K (20 mg/ mL). In-cubation occurred at 55°C with rotation. Following incubation, 250 µl phenol:chloroform:isoamyl alcohol (25:24:1) was added and samples were rotated for 10 minutes then centrifuged at 10,000 rpm for 10 minutes before transferring the aqueous phase to a new microfuge tube. This process was done twice more, first using another 250 µl phenol:chloroform:isoamyl alcohol (25:24:1), then using 250 µl chloroform:isolamyl alcohol (24:1). Two volumes of cold 100% ethanol, 1 volume of 2.5 M ammonium acetate and 1 µl glycogen were added to the final aqueous phase to precipitate the DNA. The samples were stored at -20°C for a minimum of 12 hours before three rounds of centrifugation for 10 min at 15,000 rpm at 4°C. The supernatant was subsequently discarded and the precipitate washed with 500 µl cold 70% ethanol. The precipitated DNA was left to dry at room temperature. Once dried, the DNA was resuspended in 50 µl of TE (TrisEDTA) buffer (0.01 M Tris-HCL (pH 8.0), 0.0001 M EDTA). A negative extraction control was included for every batch of samples processed (maximum of 24 per batch). DNA extractions were quantified with a Qubit 2.0 (Ther-mofisher Scientific Inc., United States) and were sent to Novogene UK or The Ramaciotti Centre for Genomics, Australia for library preparation and whole genome sequencing, aiming for 5X coverage on the Illumina Novaseq 6000 platform (Illumina, San Diego) and generating 150 bp paired-end reads.

### Identification of parasite sequences

To characterise parasitic reads from the WGS data, the approach used by Nichols *et al*. (53) was employed as follows. We used fastp to trim adapter content (10 bp) and remove low-quality base calls from the raw sequence data (60). Next, the filtered sequencing data for each sample were aligned to an existing *Z. lat-eralis* assembly (61) using the BWA-MEM algorithm (62). A host k-mer library was generated from a *Z. lateralis* assembly using the PathSeq pipeline from the Genome Analysis Toolkit (GATK) (63). The host k-mer library consists of a hash table, storing k-mers generated from consecutive positions in the reference genome sequence. A Burrows-Wheeler Aligner (BWA) image of the reference genome was also generated using the BWA-MEM index image file creator in GATK (62). Sequences from each sample were then compared to the host k-mer library using a fast k-mer search. We used the default PathSeq settings to identify reads belonging to the host: reads were removed if there was a minimum of one k-mer match to the host (63) and the BWA-MEM algorithm was employed again to align the remaining reads with the host reference image, also generated above. Reads with an identity score above 30 were discarded.

Following host sequence removal, further quality control was carried out with PathSeq (63). Reads with an excess of A/T or G/C content (29 out of 30 bases per window (64)), duplicate sequences and those shorter than 60 bp were discarded. Low-complexity sequences and low-quality bases were also masked using the sDUST algorithm (65). Read ends were further trimmed according to base quality (Q = 15).

While we considered assembling longer contiguous sequences from the remaining sequences, our ability to detect certain parasites decreased following assembly. This may be because some parasite DNA is likely to be present at extremely low levels in the host DNA and therefore will not have sufficient coverage for assembly, and because parasite reference availability is limited for longer sequences. Therefore, we opted to use shorter sequences for classification but to impose a stringent confidence filter on the subsequent taxonomic assignments. The factors to consider when making decisions about extracting parasite sequences from WGS data of the host are outlined in full in Nichols *et al*. (53).

Reference database completeness is known to influence the outcome of taxonomic assignments (66, 67). Therefore, we compiled a custom reference database in Kraken 2 to maximise parasite coverage. We utilised the pre-built bacteria, Protozoa, UniVec and human databases in Kraken 2. We then compiled additional reference sequences from various sources (Table S1). We downloaded all available genome references for apicomplexans, fornicata, cercozoans, parabasalids and euglenozoans from RefSeq (68, 69). A complete list of NCBI taxonomic sequence identifiers was generated for the same taxonomic groups using TaxonKit (70). We used Biopython to download the sequences from NCBI after removing any sequences labelled as ‘uncultured’ or ‘environmental’ (71). All genomic references from Worm-Base ParaSite (72) and the Eukaryotic Pathogen Genomics Database Resource (EuPathDB) (73) were also incorporated. Finally, the PR2 database was added following filtering to remove sequences labelled uncultured, environmental, fungal and metazoan. Kraken 2 derives a compact hash table of k-mers from these reference sequences to be queried against during sequence assignment.

Sequences were assigned using the custom parasite reference database with Kraken 2 (74). Following taxonomic assignment, we searched for information on previously recorded primary hosts and categorised the primary hosts for each parasite genus as bird, other vertebrate, invertebrate, or plant, accordingly. We removed parasites that have not previously been recorded in birds or other vertebrates. For the detected parasite genera, we collated information from the literature on their life histories which enabled assignment to the following transmission categories: flying insect vector, tick-borne, transmission involving secondary hosts other than ticks or flying insects (i.e. parasites with complex life cycles) or direct (i.e. parasites with simple life cycles). Table S2 contains a summary of the identified parasites, including available life history information.

Confidence scores for each taxonomic assignment were generated using Conifer (75). These were calculated according to the fraction of k-mers assigned to the final taxonomic identity, its descendants and ascendants, giving the Root-to-Leaf (RTL) confidence score. We applied a confidence score filter of 0.7 for the paired-end dataset. We previously found that applying a conservative confidence score to relatively short paired-end sequences resulted in fewer apparently spurious parasite identifications, i.e. fewer parasites that have not been previously reported in birds; and reasonably consistent identification of selected parasites (haemosporidians) while continuing to capture a large amount of diversity (53).

## Statistical analysis

### Is parasite diversity higher on the mainland compared to islands?

Statistical analysis was conducted in R version 4.4.2 (76). To test the island biogeography expectation of higher parasite richness in mainland compared to island populations, we used the vegan package version 2.6-10 to estimate Chao2 richness based on parasite genera detected for each population (77). To account for uneven sampling across populations, we conducted random sub-sampling with replacement and repeated the process with 1,000 bootstraps. We sub-sampled to n = 5 to reflect the lower sample sizes for some islands (see Fig. 1 for sample sizes). However, in cases where sample size was < 5, such as for Ambae (n = 3) and Vanua Lava (n = 2), Chao2 was estimated without sampling with replacement. Next, a pairwise permutation test with a Bonferroni correction for multiple comparisons was used to compare Chao2 estimates for each island population to the Australian continent. Each comparison was repeated 10,000 times.

We tested whether the Australian continent (‘the mainland’) represents a regional source for parasite communities in the island populations by looking for evidence of community nestedness. The Nestedness metric based on Overlap and Decreasing Fill (NODF) was generated using the vegan package (77). To assess whether observed nestedness differed from that expected by chance alone, we compared our matrix to 999 simulations of a row-constrained null model, in which parasite richness on each island (row totals) was conserved across the matrix to account for uneven sampling size, while parasite incidence (column totals) was randomised. We performed the analyses on two datasets: in the first, we included the mainland, and each island population individually; and in the second, we combined data from islands within each of the French Polynesia and Southern Melanesia archipelagos. We then considered each archipelago as single units for analysis, to reflect the geographical clustering of the islands.

To examine how variation in parasite communities differed in host individuals from the islands compared to the Australian continent, we implemented an extreme gradient boosting (XGBoost) classification algorithm using the XGBoost package 1.7.11.1 (78). We aimed to test whether a host individual’s origin (mainland or island) could be predicted according to a ‘parasite profile’. The parasite profiles consisted of the number of genera detected in each host individual per transmission category. The predicted variable was host origin (i.e. island or mainland). We subsequently extracted importance scores for each of the four transmission categories in classifying host individuals as mainland or island origin. Although XGBoost has been shown to work effectively on imbalanced datasets in the past (79), it is nonetheless likely to cause a decline in model performance if one class is overrepresented (80). Therefore, we applied class weighting to the model to account for the imbalanced sample sizes in the island (n = 329) and mainland (n = 60) categories. Specifically, we used the ratio of mainland to island host individuals to adjust the gradient updates during training by increasing the weight of errors made on the underrepresented class (mainland hosts), thereby preventing the model from being biased toward the majority class (island hosts). We examined the distribution of probabilities output by the model to confirm that class imbalance was not causing bias in predictions (Figure S1). We used leave-one-out cross-validation (LOOCV), where one host individual was withheld in each iteration of the model. We calculated Area Under the Receiver Operating Characteristic Curve (AUC ROC) scores to assess the model.

To further illustrate the output of XGBoost, we tested the difference in mean number of parasite genera of each transmission category detected per host. We utilised a two-way ANOVA using host origin and transmission category as factors, followed by a post-hoc Tukey HSD test to explore pairwise comparisons. We also tested differences between the prevalence per transmission category (i.e. pooling parasite genera into four transmission categories for island and mainland populations) on the island and mainland using four independent Fisher’s Exact Tests (one per transmission category) and applied a Bonferroni correction to control for multiple comparisons. The adjusted threshold significance was *α* =0.0125.

### Is parasite richness higher on less isolated, larger islands?

To test whether parasite richness increased with island size and decreased with island isolation, we fitted generalized linear models (GLMs) to compare Chao2 richness among islands. For each model, we used the same set of predictor variables: island area, island isolation, and latitude. To calculate island area and isolation, we utilised the Natural Earth 10m landmass data (81, 82). Areas were computed on the WGS84 ellipsoid (EPSG:4326). We considered two metrics of isolation to capture larger-scale and regional-level isolation. First, we calculated the distance to mainland as the shortest possible distance between the coast of mainland Australia and each island (km). Second, we followed the procedure outlined by Itescu et al. (83) and Weigelt and Kreft (84), generating a concentric 100 km buffer around each island and calculating the area of landmass it contained, excluding the island itself. This captures a more regional isolation level, which is relevant when a dataset comprises both single isolated islands and islands that fall within archipelagos. Isolation metrics and area varied across several orders of magnitude. To reduce positive skew, we applied a log_10_ transformation to island area and both measures of isolation prior to modelling. To control for the possibility that parasite communities vary according to latitude (10, 85), latitude was included as a covariate in the models. Our sampling took place over two decades, over which time parasite communities may be liable to change (41). However, we were unable to include sampling year as a random effect because most islands were sampled in a single year, therefore including year as a random factor would prevent models from converging. The models were also weighted to account for uneven sample sizes among populations.

To avoid inclusion of collinear predictor variables in the models, pairwise Pearson correlation coefficients were calculated between each predictor variable to ensure that none exceeded a correlation threshold of r > 0.7, which they did not. Variance inflation factors (VIFs) were also computed using the car package to quantify multicollinearity (86). The moderate correlation between proximate island isolation and island area led to elevated VIF scores (Table S3). Therefore, we excluded regional isolation and recalculated VIF scores. All scores were below two, indicating acceptable levels of collinearity for inclusion in the model.

In the first GLM, we used the log_10_-transformed mean estimates for Chao2 richness per island as the response variable and specified a Gaussian distribution of errors. In the second GLM, we classified parasite genera into direct, flying insect vector or other secondary host transmitted parasites and calculated Chao2 richness for each category. We excluded tick-borne parasites from this analysis as the overall richness detected was low (only two genera were detected), and Chao2 estimates were thus unlikely to be informative (87). We found that using a log_10_-transformation was not sufficient to adjust the right-skewed data, so we instead used a gamma distribution of errors to accommodate the resultant non-negative, count-like richness measures. Despite being count-like, Chao2 is in fact a bias-corrected estimate derived from incidence data (88). Therefore, Chao2 estimates are continuous, non-integer values which cannot be modelled using error distributions that are commonly used for count data. Following model fitting, we used the DHARMa package (89) to assess model fit (Figure S2).

For both GLMs, we performed model selection and averaging using the MuMIn package (90). All candidate models were compared based on Akaike’s Information Criterion corrected for small sample sizes (AICc). Models within ΔAICc< 2 of the top-ranked model were considered equally plausible, and we generated model-averaged predictions across this top model set. Alternatively, if no other models were within ΔAICc < 2, we selected the top model. Model selection tables are included in Tables S5 and S6.

## Results

We characterised 48 parasite genera from 384 individual silvereye whole genome sequences. Of these, the majority require a secondary host for transmission (45% of total genera detected), followed by equal numbers of directly transmitted genera (26%) and those transmitted by flying insect vectors (26%), and lastly by tick-borne parasite genera (4%). Of the four groups, flying insect vector-transmitted parasites had the highest prevalence, with at least one genus detected in every population.

### Is parasite richness higher on the mainland compared to islands?

Parasite richness was generally higher on the mainland than for island populations, with 24 of the 25 island populations having significantly lower Chao2 richness estimates (Fig. 1). The exception was the Heron Island population, with a significantly higher, though highly variable estimated Chao2 compared to the mainland.

The parasite communities of host populations exhibited significant nestedness (Fig. 2). In the analysis considering each island population separately, the NODF was 63.79, substantially higher than expected under the null model (null model mean = 47.31, 95% CI = 43.07-51.42, p < 0.001). This corresponds to a standardized effect size of 7.76. The mainland fell at the apex of the nestedness structure, with island parasite communities forming subsets of the genera found on the mainland (Fig. 2A). In the analysis that considered islands within Southern Melanesia and French Polynesia at the archipelago level, the observed NODF was 71.68 (null model: mean = 59.432, 95% CI = 53.03-65.55, p < 0.001) and the standardised effect size was 3.87. However, in this analysis, Southern Melanesia fell at the apex of the nestedness structure (Fig. 2B).

**Fig. 2.**
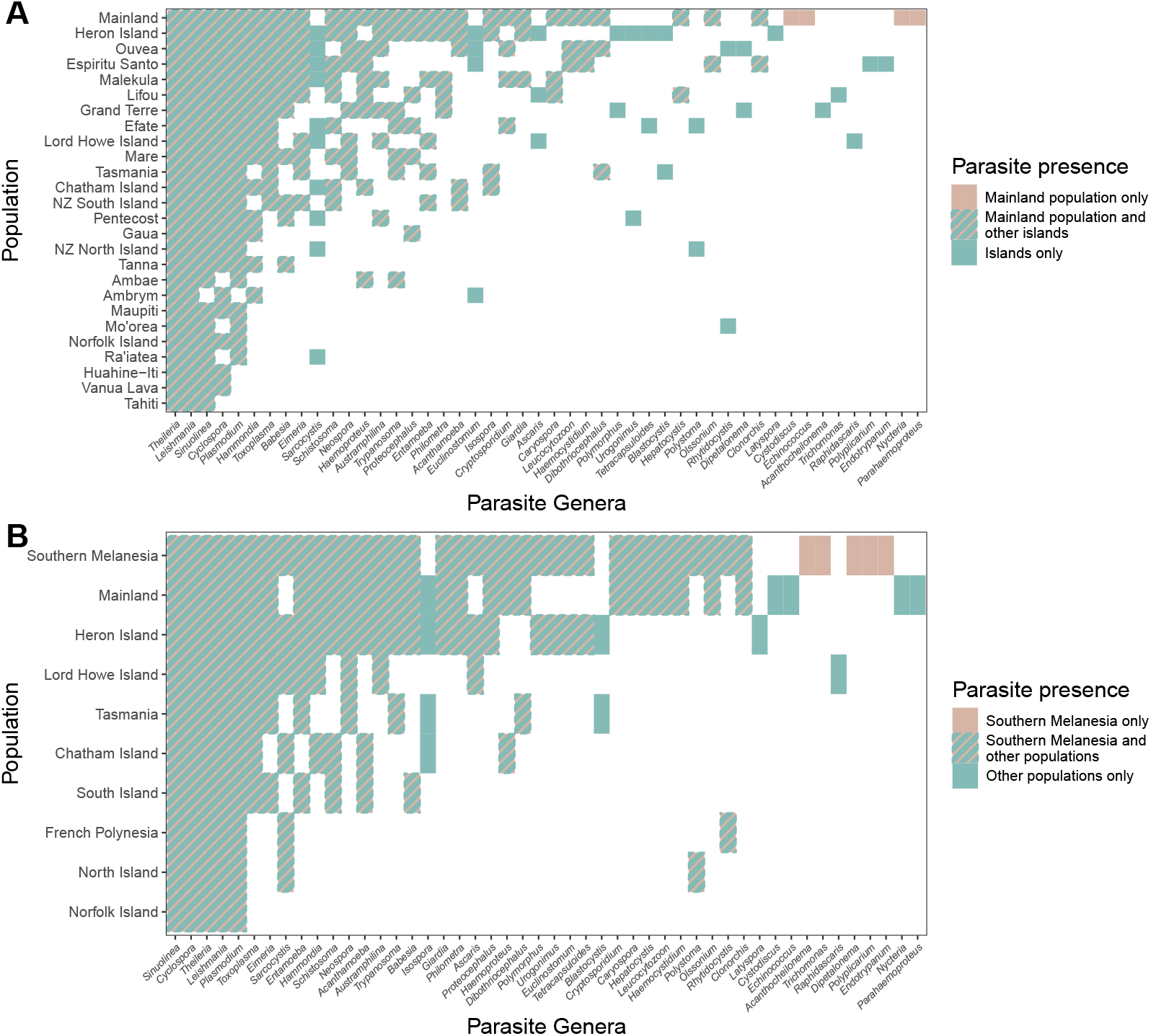
Nestedness structure amongst A) the mainland and each island population and B) the mainland, single islands, and the Southern Melanesia and French Polynesia archipelagos (grouping islands within each of these archipelagos).

According to LOOCV, the XGBoost model had a classification accuracy of 82% and an AUC ROC score of 86%, indicating that the parasite profile, i.e. the number of parasite genera detected per individual host per transmission category, provides discriminatory power to identify host samples sourced from an island versus the mainland. Feature importance scores extracted from the model indicated that flying insect vector-transmitted parasites were most informative in this distinction (Fig. 3A). Furthermore, the mean number of genera detected per individual host was significantly higher for mainland than island populations for direct (ANOVA, mean difference = 0.41 95% CI = 0.09-0.73, p < 0.001) and flying insect vector-transmitted parasites (ANOVA, mean difference = 1.36, 95% CI = 1.06-1.67, p < 0.001) whereas there was no significant difference for parasites transmitted by ticks (ANOVA, mean difference = 0.04, 95% CI = -0.27-0.35, p = 0.999) or other secondary hosts (ANOVA, mean difference = 0.08, 95% CI = -0.29-0.46, p = 0.997; Fig. 3B).The prevalences of directly transmitted (Fisher’s Exact Test, Odds Ratio (OR) = 9.07, 95% CI = 2.30-78.51, Bonferroni-adjusted p < 0.001), other secondary host transmitted (OR = 4.93, 95% CI = 2.46-10.48, Bonferroni-adjusted p < 0.001), and tick-borne (OR =∞, 95% CI = 3.65-∞, Bonferroni-adjusted p < 0.001) parasites were higher on the mainland than in the island populations. No difference was found for flying insect vector-transmitted parasites between island and mainland populations (OR =∞, 95% CI = 0.44-∞, Bonferroni-adjusted p = 1.00).

**Fig. 3.**
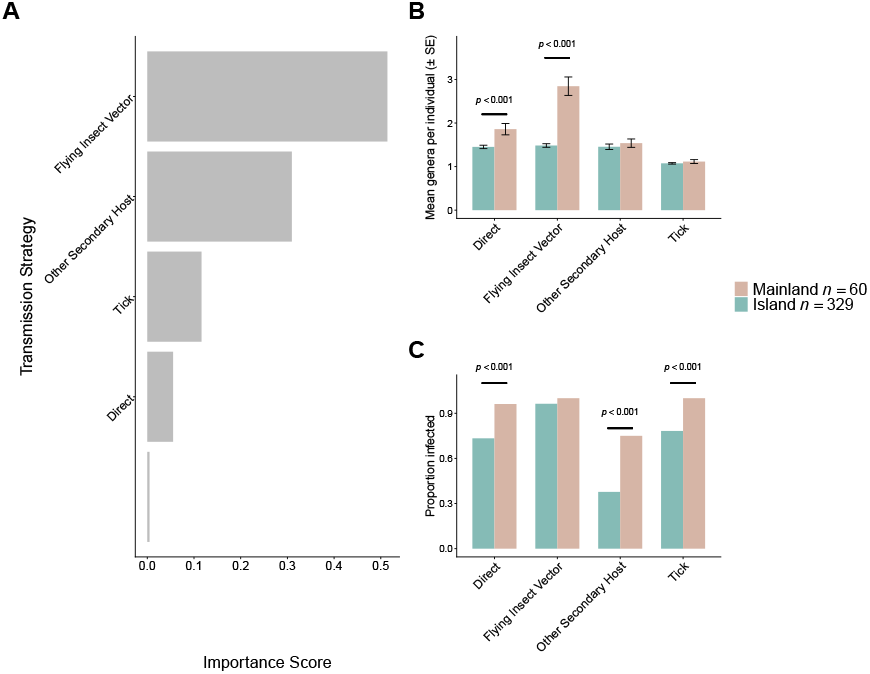
A) Importance scores extracted from XGBoost when classifying 384 individual hosts into island and mainland origins according to their parasite profile (number of parasite genera per transmission category detected per individual host) B) mean number of genera of parasites detected per individual host according to transmission category, p values generated using an ANOVA and Tukey HSD test and C) proportion of individual hosts sampled from the mainland and island populations with parasites detected according to transmission category, p values derived from Fisher Exact Test with Bonferroni correction.

### Is parasite richness higher on less isolated, larger islands?

For total Chao2 richness per island, two models were averaged based on AICc (Tab. 1). Island area was included in one of the two models, and because the models were similarly weighted, island area was attributed a weighted importance score of 0.49, while latitude and isolation from the Australian continent were attributed weighted importance scores of 1. While there was support for including all three variables in the models, isolation from the Australian continent had the most substantial effect size, with richness decreasing as isolation from the Australian continent increased (Fig. 4A). Although island area and latitude did explain some of the variation in the data, it was not possible to discern the direction of the relationships.

**Table 1.**
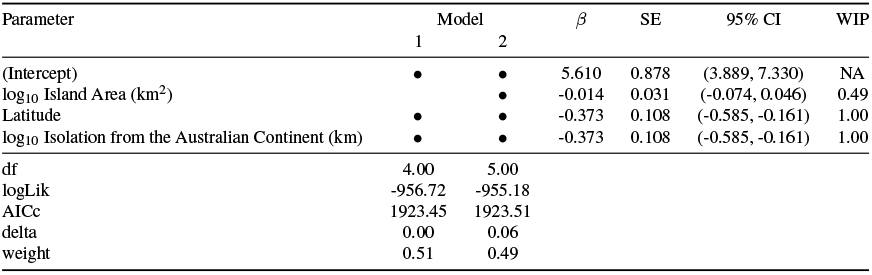
Model-averaged estimates for GLM predicting total Chao2 richness based on parasite genera detected per island population. Estimates, standard errors, 95% CI and weighted importance scores (WIP) are based on model averaging across the top two models (ΔAICc < 2; Table S4). The parameters included in each model (1 and 2) are indicated by black dots; for each model in the top set, we also present degrees of freedom (df), Log-likelihood (logLik), the difference in AICc from the top model (delta) and the relative support for each model according to the AICc scores (weight).

**Fig. 4.**
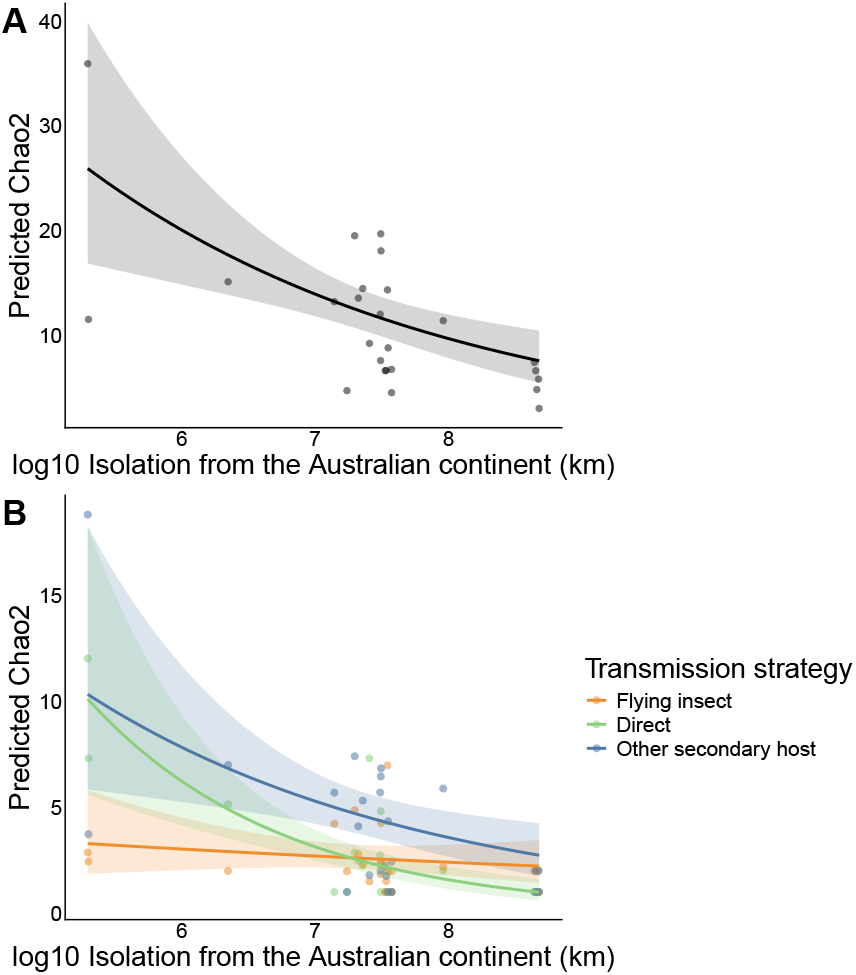
Predicted Chao2 richness as a function of isolation from the Australian continent for A) total Chao2 richness per island and B) per transmission strategy. Predictions are based on an average of the top two models with < 2 AICc (A) or the top GLM according to AICc values (B). Points represent observed island population values used to generate predictions.

For Chao2 richness per transmission category on each island, a single top model was selected based on AICc scores (Table S4). The top model included every variable, although it was not possible to discern the direction of the relationship between Chao2 richness and island area, isolation from the Australian continent, and latitude, as the 95% CI encompassed zero (Tab. 2). However, we found substantial variation between Chao2 richness for each transmission category: with directly transmitted parasites having the overall highest richness, followed by parasites transmitted by other secondary hosts and finally, flying insect vector-transmitted parasites. We also found that the relationship between Chao2 richness and isolation from the Australian continent was dependent on transmission strategy (Fig. 4B), with distance having almost no impact on parasites that rely on flying insect vectors but a substantial decline in richness for parasites utilising direct transmission strategies and those involving other secondary hosts. Tab. 3 summarises the results obtained from each of the statistical tests performed.

**Table 2.**
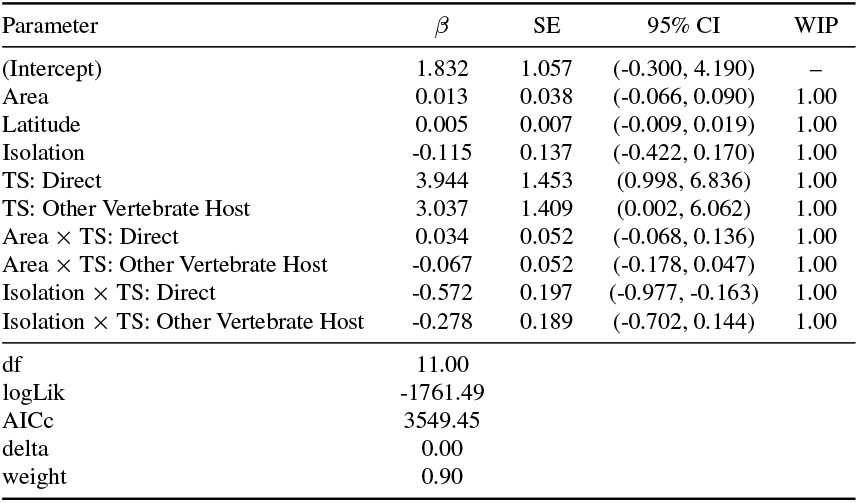
Model estimates from the top GLM predicting Chao2 richness per transmission category on each of 25 islands. Estimates, standard errors, 95% CI and weighted importance scores (WIP) are based on the top model (ΔAICc < 2; Table S5). The parameters included in each model are indicated by black dots; for each model in the top set, we also present degrees of freedom (df), Log-likelihood (log-Lik), the difference in AICc from the top model (delta) and the relative support for each model according to the AICc scores (weight). Area = log_10_ Island Area (km^2^), Isolation = log_10_ isolation from the Australian continent (km), TS = Transmission Strategy.

**Table 3.** Summary of results obtained from each comparison made (island vs mainland or island vs island) for different groups of parasites.

| Comparison and parasite group | Parasite measure | Direction |
| --- | --- | --- |
| **Island vs mainland** |  |  |
| Total | Richness (Chao2) | Higher on mainland than islands |
|  | Nestedness | Islands nested within mainland |
| Direct (simple) | Richness (count) | Higher on mainland than islands |
|  | Prevalence | Higher on mainland than islands |
| Flying insect vector (complex) | Richness (count) | Higher on mainland than islands |
|  | Prevalence | No difference |
| Other Secondary Host (complex) | Richness (count) | No difference |
|  | Prevalence | Higher on mainland than islands |
| Tick (complex) | Richness (count) | No difference |
|  | Prevalence | Higher on mainland than islands |
| **Island vs Island (isolation)** |  |  |
| Total | Richness (Chao2) | Declines with increasing isolation |
| Direct (simple) | Richness (Chao2) | Declines with increasing isolation |
| Flying insect vector (complex) | Richness (Chao2) | No difference |
| Other Secondary Host (complex) | Richness (Chao2) | Declines with increasing isolation |
| **Island vs Island (area)** |  |  |
| Total | Richness (Chao2) | No difference |
| Direct (simple) | Richness (Chao2) | No difference |
| Flying insect vector (complex) | Richness (Chao2) | No difference |
| Other Secondary Host (complex) | Richness (Chao2) | No difference |

## Discussion

By extracting parasite DNA from low-coverage whole genome sequences of hosts, we empirically demonstrate how parasite distributions reflect the expected patterns of island biogeography. Notably, we found that distribution patterns were differentially driven by parasites utilising varied transmission strategies. Overall, parasite species richness was higher on mainland Australia compared to island populations and, among island populations, parasite richness decreased with distance from the continental mainland. Taken together with our nestedness analysis, it appears that parasite diversity in the region is sourced from both the Australian continent and Melanesian archipelagos.

### Isolation and parasite life history traits influence parasite richness and prevalence

Our results clearly demonstrate that transmission strategies influence parasite patterns of island biogeography. For directly transmitted, simple life cycle parasites, we found lower parasite richness in island populations, with richness declining with distance from the continental mainland. These results indicate that when the primary dispersal mechanism is co-dispersal with the host, island isolation acts as a dispersal filter simultaneously for both host and parasite. Although studies which test the rules of island biogeography in directly transmitted parasites are rare, reduced richness of directly transmitted helminths with distance from the nearest island has previously been shown for fish parasites within the Pacific Line Islands (25) and is also demonstrated by reduced haplotype diversity with distance from the mainland in the nematode parasite, *Heligmosomoides polygyrus* infecting mice in the Mediterranean (91). Parasites with complex life cycles showed varied patterns of island biogeography. Some variation was expected given constraints associated with life cycle complexity, however other results highlight the need for additional ecological and evolutionary considerations. Parasites that utilise intermediate hosts are expected to be more limited in their establishment success than directly transmitted parasites (40, 41), and this should lead to stronger patterns of island biogeography. However, we detected no significant difference overall in richness between island versus mainland populations for such parasites, and although richness decreased with increasing island isolation, the effect was weaker than for directly transmitted parasites. A possible explanation for these patterns is that utilisation of a paratenic host (a host that can harbour the parasite in a dormant state) may facilitate persistence in newly colonised environments as parasites do not require immediate availability of definitive hosts (38, 92, 93). Therefore, rather than being a limitation, intermediate host use could enhance opportunities for colonisation and persistence, thereby weakening richness and distance effects.

Parasites transmitted by flying insect vectors exhibited a reduction in genus richness on islands compared to the mainland, yet this richness did not decline further with increasing island isolation. The overall reduction in richness on islands followed expectations as successful transmission requires a suitable vertebrate host and compatible flying insect vector (94). Variation in the dispersal propensities and flight mechanisms of flying insects (95), combined with the relatively limited availability of habitat types on islands (32, 96) may restrict diversity of vectors able to establish, thereby constraining the diversity of vector-dependent parasites (97). However, the observed patterns suggest that for the subset of parasites whose vectors can traverse long distances, island isolation likely exerts a weaker influence on colonisation success. For example, wind-assisted dispersal of *Anopheles* mosquitos can promote travel over 300 km in a 9 hour flight period (98). This finding has important implications for future disease projections under climate change. In particular, the dispersal of flying insect vectors may be augmented globally due to lower air densities resulting from warming temperatures (99, 100) and changing wind patterns (101). Consequently, highly dispersive flying insect vectors may be increasingly able colonise remote geographic reaches, thereby creating new opportunities for transmission.

In contrast to parasite richness, prevalence of flying insect vector-transmitted parasites remained similarly high on the mainland and the islands. A pattern of high prevalence of island parasites has been reported previously (102–104) and reflects a well-established biogeographic trend for free-living organisms whereby the subset of species that occur on islands often reach higher population densities (105). Maintenance of comparable parasite prevalence on islands (cf mainland) may therefore be supported by high host densities, which facilitate frequent transmission opportunities (106, 107).

### Island area and parasite richness

Although area was important in explaining differences in parasite richness among islands, we did not detect a positive correlation between richness and island area, as expected under the rules of island biogeography. While our study included some very large islands, e.g. the North and South Islands of New Zealand, which are*∼* 114,000 and *∼* 150,000 km^2^, respectively, most of the islands included would be considered small according to the UNESCO definition (< 2000 km^2^; Falkland and Custodio (108)). The lack of a positive correlation between richness and island area may reflect the Small Island Effect (SIE), which describes how the relationship between species richness and island area can be decoupled for small islands (109). The mechanisms underlying this effect include reduced habitat diversity on small islands (110, 111), edge effects (109), or disequilibrium between colonisation and extinction events. Therefore, for small islands, stochastic processes and ecological factors may have a greater impact on richness than area-dependent relationships predicted by classical island biogeography.

While we do not know which explanations for SIEs may be at play in this system, one island in our dataset clearly exemplifies the decoupling described by the effect. Despite being the smallest island in our dataset, Heron Island (0.29 km^2^) hosted the richest parasite community. The proximity of Heron Island to the mainland (*∼*80km) likely bolsters the parasite community via relatively frequent arrival of a wide variety of avian vagrant species (112) and vectors e.g. the arrival of vectors following episodes of offshore winds was proposed to explain appearance of *Plasmodium* infection in the silvereye population on Heron Island (37). The position of Heron Island immediately below mainland Australia in the nestedness analysis supports the idea that parasites readily colonise from the mainland. The high parasite richness of Heron Island silvereyes may be further elevated due to sharing of parasites from seabirds. Several parasite genera detected on Heron Island were not detected on the Australian mainland, some known to parasitise seabirds e.g. *Sarcocystis* (113, 114), *Euclinostomum* (115, 116) and *Polymorphus* (117, 118). Heron Island supports high-density breeding populations of Black Noddies (*Anous minutus*) and Wedge-tailed Shearwaters (*Ardenna pacifica*) (119, 120). These produce substantial amounts of guano e.g. estimates of over 100 tonnes per annum for Black Noddies alone on Heron Island (121). The abundance of guano likely provides opportunities for parasite transmission as the silvereyes routinely forage in Black Noddy nests, as well as consume food items that have been regurgitated by noddies. However, we recognise that the distribution of Heron Island Chao2 estimates is highly variable, and it is likely that further sequencing would be required to fully characterise the parasite profile of the silvereyes in this population to better understand the drivers of the high average Chao2 estimates.

### Sources of parasite fauna

The nestedness analysis identified the Australian mainland as the dominant source of parasites for the sampled island populations. However, the detection of several parasite genera on the islands but not on the Australian mainland indicates that multiple source populations may be involved in determining regional parasite assemblages. When analysed at the archipelago scale, Southern Melanesia occurred at the apex of the hierarchical structure of nestedness. New Guinea is a major source for much of the regional avifauna in Southern Melanesia (122–125), and it follows that movements of birds from New Guinea have provided a source of parasite diversity in Southern Melanesia and beyond.

Host colonisation history likely further influences the parasite assemblages detected across islands. Silvereyes have colonised islands throughout the South Pacific over a range of timescales. Some populations in Southern Melanesia are estimated to have colonised hundreds of thousands of years ago, while others are the product of recent, human-mediated introductions. For instance, silvereyes in French Polynesia exhibit low parasite richness. These populations were established from an introduction of a small number of silvereyes from South Island New Zealand in the 1930s (126, 127) and the low parasite richness observed may reflect failure to colonise with the host, or early loss post-colonisation (MacLeod et al. 2010). In addition, parasite gain from other avifauna must be minimal. The native avifauna may have had historically low parasite diversity due to the highly isolated nature of the archipelago. However non-native introductions that occurred in the late 19th and early 20th centuries (128) conceivably provide opportunities for parasite sharing and with time, new host-parasite relationships in French Polynesian silvereyes may arise.

### Caveats

The limited understanding of the breadth of host associations and patterns of host use in our system makes it challenging to empirically assess how host specificity influences parasite distributions across islands. A previous study found that host specificity of parasites significantly decreased on islands when comparing the outcomes of 28 studies on 16 species of mammal (107). Therefore, it is likely that being a host generalist confers a benefit to parasites colonising islands. However, there is limited information on the factors that shape host specificity, and levels of specificity are liable to change with environmental variability (129). Furthermore, parasite specificity is likely to vary with transmission strategy. For example, the transmission of tick-borne parasites will be further complicated as ticks themselves have complex life cycles (130, 131) and display varying levels of host specificity (132). There has been a sustained interest in characterising the degree of host specificity for some avian parasites (133–137). Such work may help to elucidate the impact of host specialisation on island colonisation in the future.

Re-purposing Whole Genome Sequence data to characterise affiliated parasites involves some trade-offs. While the approach enables detection of a broad range of parasites at the genus level, some diversity may be overlooked due to the limited availability of published reference sequences. Furthermore, the lack of longer or complete genome reference sequences means that assembly of short reads into longer contiguous sequences yields fewer matches. Yet the use of short sequences introduces the possibility of assignment to an incorrect reference. To combat these collective issues, we used short sequences rather than assembling longer sequences, but applied a stringent confidence filter for identifications, and classified only to the genus level (53). Our approach reflects the decision to maximise diversity capture, despite introducing some false-positive identifications.

## Conclusion

By leveraging existing WGS data from an island-colonising bird, we provide evidence that large-scale parasite distributions follow the rules of island biogeography, but these distributions are mediated by aspects of parasite life history. This suggests that inconsistent biogeographical patterns revealed in previous studies that focus on particular parasite groups e.g. (22, 102, 138) can be at least partially explained by variation in life history, an insight that is only revealed by examining a broad range of parasites. While the increasing availability of parasite sequence data in public databases will enhance analyses of distribution patterns, a concerted effort is also needed to gather information on parasite ecology and life history traits (40). Such integrative data will enable a more nuanced understanding of parasite distributions over space and improve forecasts of future distributional changes, including the spread of disease agents.

## ACKNOWLEDGEMENTS

We are grateful to the many people who facilitated this work. We thank the chiefs and landholders of Vanuatu and New Caledonia for granting access to field sites. We are grateful to those who assisted with fieldwork and/or logistics for fieldwork: N. Clark, R. Black, D. Treby, J. LeBreton, F. Cugny, W. Waheoneme, O. Hebert and A. Rouquie, G. Kakue (New Caledonia); F. Robertson and C. Sendell-Price (Heron Island); S. Geiger, O. Boissier, R. Hills, S. Totterman, O. Drew, K. Ser, W. Ser, J. Saksak, A. Phillimore (Vanuatu), I.P.F. Owens, N. Clark, F. Robertson, M. Massaro, D. Potvin (Australian mainland); J. Kikkawa (Norfolk Island); I.P.F. Owens (Lord Howe Island); P. Park, A. Fletcher, P. Gray, F. Robertson (Tasmania); P. Schweigman, D. Onely, F. Robertson (New Zealand); M. Bell (Chatham Island); N. Davies, O. Grant, C. Sendell-Price (Moorea, French Polynesia). We also thank D. Bass and M. Kamouyiaros for advice and discussions about the bioinformatics required to extract parasite sequences from the data.

The work was conducted under permits from the Direction de l’Environnement Province Sud and Direction Du Développement Economique (New Caledonia and Loyalty Islands with thanks to G. Kakue); Vanuatu Environment Unit letters of permission and permits provided by E. Bani and we further thank D. Kalfatak and T. Tiwok for their assistance (Vanuatu); Délégations régionales à la recherche et à la technologie (French Polynesia); Lord Howe Island Board; Environment Australia (Norfolk Island); Queensland Department of Environment and Resource Management; Parks and Wildlife Service (New South Wales with thanks to M. Massaro); Parks and Wildlife Service (Tasmania); New Zealand Department of Conservation Te Papa Atawhai (New Zealand); Australian Bird and Bat Banding Scheme project (SMC), and New Zealand National Banding Office (BCR).

## Animal studies

Ethics clearances were provided by University of Queensland ethics committee (ZOO/165/94/ARC, ZOO/520/96/ARC/PHD, ZOO/520/97/ARC/PHD), Griffith University ethics committee (ENV/01/12/AEC, ENV/07/16/AEC, ENV/06/20/AEC, ENV/24/13/AEC) to SMC, Charles Sturt University AEC to BCR and M. Massaro and Otago University AEC to BCR.

## Funding

We thank the funders of this work: the Department of Zoology, Oxford start-up and OUP John Fell Fund grant to SMC to support sequencing; Marsden Fund New Zealand to BCR and SMC to support fieldwork on mainland Australia and Heron Island and sequencing; National Geographic Society Committee for Research and Exploration Grant (9383-13) to SMC to support fieldwork in New Caledonia; Natural Environment Research Council (NERC) postdoctoral fellowship to SMC to support fieldwork in Vanuatu and New Caledonia; Percy Sladen Memorial fund to SMC to support fieldwork in French Polynesia; SN was supported by a NERC studentship (NE/S007474/1); AE was supported by a NERC studentship (NE/S007474/1) and a St John’s College Graduate Scholarship.

## Data Accessibility Statement

Raw sequence reads will be deposited in the SRA (BioProject ID to be provided upon acceptance). The associated sample metadata will also be stored in the SRA.

## Benefits Generated

Benefits from this research accrue from the sharing of our data and approach on public databases as described above.

## Author contributions

Conceptualisation: SN, SMC and BO designed the research. Investigation: SMC conducted fieldwork in Southern Melanesia, French Polynesia, New Zealand, Australia (including Tasmania, Norfolk Island, Lord Howe Island, and Heron Island); BCR and FR conducted fieldwork in New Zealand and Australia (including Tasmania); AE and FR conducted the lab work. Formal analysis: SN analysed the data and visualised the data. Supervision: SMC and BO provided supervisory oversight. Writing, original draft: SN wrote the paper with BO and SMC. Writing, review & editing: All authors provided feedback on the manuscript during the drafting process.

## Notes

### Competing Interest Statement

The authors have declared no competing interest.

