## Supplementary Material for "Transmission strategy modulates parasite biogeography in an island-colonising bird"

**Table S1.** Summary of reference sequences included in the custom Kraken 2 database.

| Source | Details | Sequences obtained (n) |
| --- | --- | --- |
| NCBI Genomes | Downloaded via the NCBI Datasets tool, including RefSeq for: | |
|  | [Apicomplexans](https://www.ncbi.nlm.nih.gov/datasets/genome/?taxon=5794) | 372 |
|  | [Cercozoans](https://www.ncbi.nlm.nih.gov/datasets/genome/?taxon=136419) | 5 |
|  | [Euglenozoa](https://www.ncbi.nlm.nih.gov/datasets/genome/?taxon=33682) | 188 |
|  | [Fornicata](https://www.ncbi.nlm.nih.gov/datasets/genome/?taxon=207245) | 36 |
|  | [Parabasalids](https://www.ncbi.nlm.nih.gov/datasets/genome/?taxon=5719) | 10 |
| [WormBase ParaSite](https://parasite.wormbase.org/ftp.html) | Version: WBPS18 | 4,856,896 |
| [EuPathDB](https://veupathdb.org/veupathdb/app/fasta-tool/genomic-sequence) | Release 64 12 Jul 2023 | 281,024 |
| [PR^2^ database](https://pr2-database.org/) | Release 5.0.0 with metazoan, fungal and uncultured sequences removed. | 101,252 |
| [NCBI Nucleotide database](https://www.ncbi.nlm.nih.gov/nuccore) | All available sequences for apicomplexans, cercozoans, euglenozoa, fornicata and parabasalids (See supplementary material for TaxIDs). | 37,153 |

**Table S2.** Parasite genera, transmission strategy and recorded host identified from 381 individual silvereye whole genome sequences.

| Parasite genera | Transmission strategy | Host | Number detected |
| --- | --- | --- | --- |
| *Acanthamoeba* | Direct | Birds | 6 |
| *Acanthocheilonema* | Flying Insect | Birds | 2 |
| *Ascaris* | Direct | Birds | 3 |
| *Austramphilina* | Other Secondary Host | Vertebrates | 11 |
| *Babesia* | Tick | Birds | 29 |
| *Blastocystis* | Direct | Birds | 2 |
| *Caryospora* | Direct | Birds | 4 |
| *Clonorchis* | Other Secondary Host | Vertebrates | 2 |
| *Cryptosporidium* | Direct | Birds | 7 |
| *Cyclospora* | Direct | Birds | 113 |
| *Cystodiscus* | Other Secondary Host | Vertebrates | 1 |
| *Dibothriocephalus* | Other Secondary Host | Birds | 5 |
| *Dipetalonema* | Flying Insect | Vertebrates | 4 |
| *Echinococcus* | Other Secondary Host | Vertebrates | 1 |
| *Eimeria* | Direct | Birds | 36 |
| *Endotrypanum* | Flying Insect | Vertebrates | 1 |
| *Entamoeba* | Direct | Birds | 7 |
| *Euclinostomum* | Other Secondary Host | Birds | 5 |
| *Giardia* | Direct | Birds | 5 |
| *Haemocystidium* | Flying Insect | Vertebrates | 12 |
| *Haemoproteus* | Flying Insect | Birds | 44 |
| *Hammondia* | Other Secondary Host | Vertebrates | 113 |
| *Hepatocystis* | Flying Insect | Vertebrates | 3 |
| *Isospora* | Other Secondary Host | Birds | 8 |
| *Latyspora* | Other Secondary Host | Vertebrates | 1 |
| *Leishmania* | Flying Insect | Birds | 355 |
| *Leucocytozoon* | Flying Insect | Birds | 14 |
| *Neospora* | Other Secondary Host | Birds | 11 |
| *Nycteria* | Flying Insect | Vertebrates | 3 |
| *Olssonium* | unknown | Vertebrates | 3 |
| *Parahaemoproteus* | Flying Insect | Birds | 4 |
| *Philometra* | Other Secondary Host | Vertebrates | 9 |
| *Plasmodium* | Flying Insect | Birds | 142 |
| *Polymorphus* | Other Secondary Host | Birds | 2 |
| *Polyplicarium* | Direct | Vertebrates | 1 |
| *Polystoma* | Direct | Vertebrates | 2 |
| *Proteocephalus* | Other Secondary Host | Vertebrates | 6 |
| *Raphidascaris* | Other Secondary Host | Vertebrates | 1 |
| *Rhytidocystis* | Other Secondary Host | Vertebrates | 2 |
| *Sarcocystis* | Other Secondary Host | Birds | 12 |
| *Schistosoma* | Other Secondary Host | Birds | 17 |
| *Sinuolinea* | Other Secondary Host | Vertebrates | 242 |
| *Tetracapsuloides* | Other Secondary Host | Vertebrates | 2 |
| *Theileria* | Tick | Vertebrates | 303 |
| *Toxoplasma* | Other Secondary Host | Birds | 62 |
| *Trichomonas* | Direct | Birds | 1 |
| *Trypanosoma* | Flying Insect | Birds | 17 |
| *Urogonimus* | Other Secondary Host | Birds | 3 |

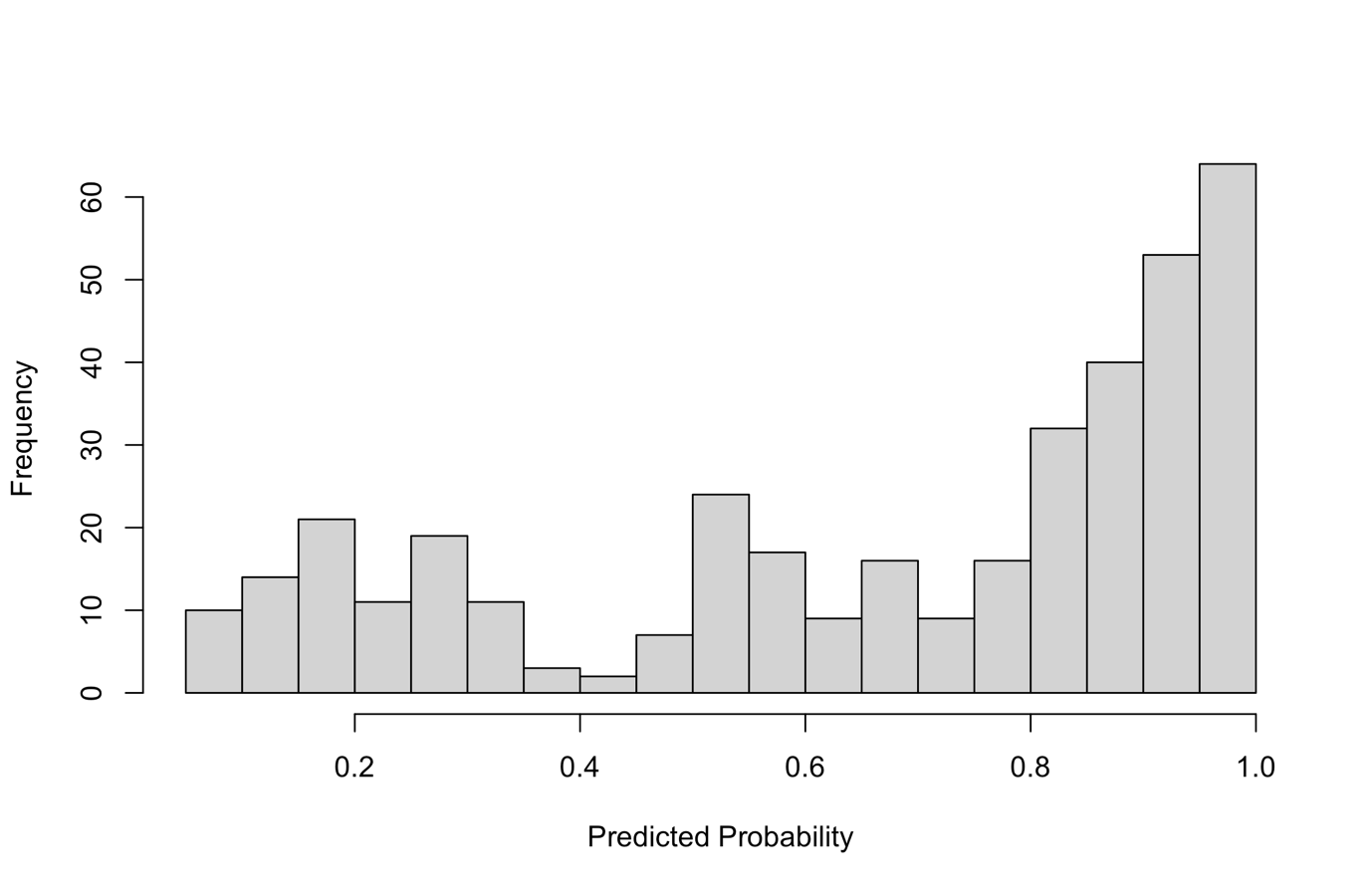

**Figure S1.** Distribution of predicted probabilities output from XGBoost when classifying 381 individual silvereye into island and mainland origin groups according to their parasite profile (number of parasite genera per transmission category detected per individual host).

**Table S3.** Pairwise Variance inflation factors (VIFs) and Pearson correlation coefficients calculated for each predictor variable included in the GLMs.

| Variable | VIF (excluding Proximate Isolation**)** | VIF | log10 Island Area (km^2^) | log10 isolation from the Australian continent (km) | Latitude | Population Size |
| --- | --- | --- | --- | --- | --- | --- |
| log10 Island Area (km^2^) | 1.41 | 4.17 | 0.66 | -0.09 | -0.43 | 0.36 |
| log10 Regional Island Isolation (km^2^) | NA | 3.20 |  | 0.24 | 0.13 | -0.03 |
| log10 Isolation from the Australian continent (km) | 1.56 | 1.57 |  |  | 0.40 | -0.51 |
| Latitude | 1.47 | 2.12 |  |  |  | -0.33 |
| Population Size | 1.56 | 1.69 |  |  |  |  |

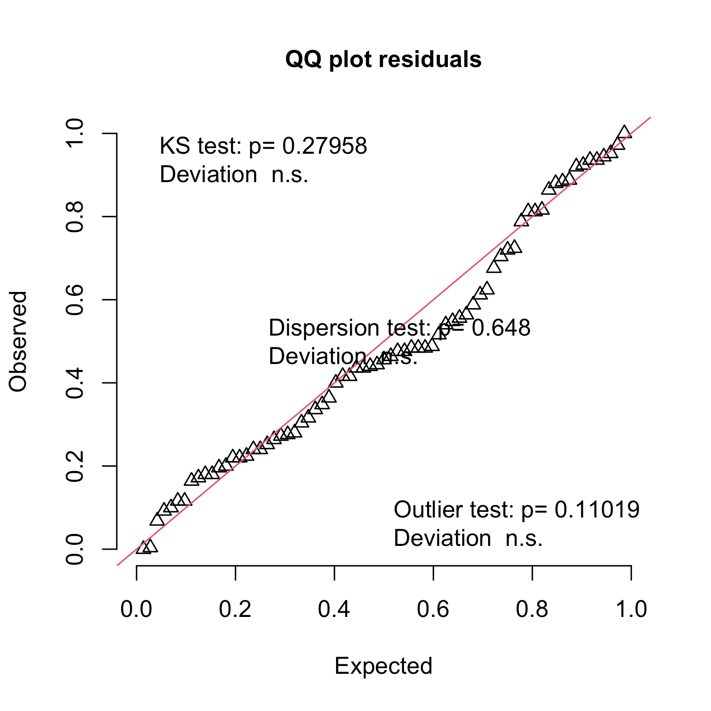

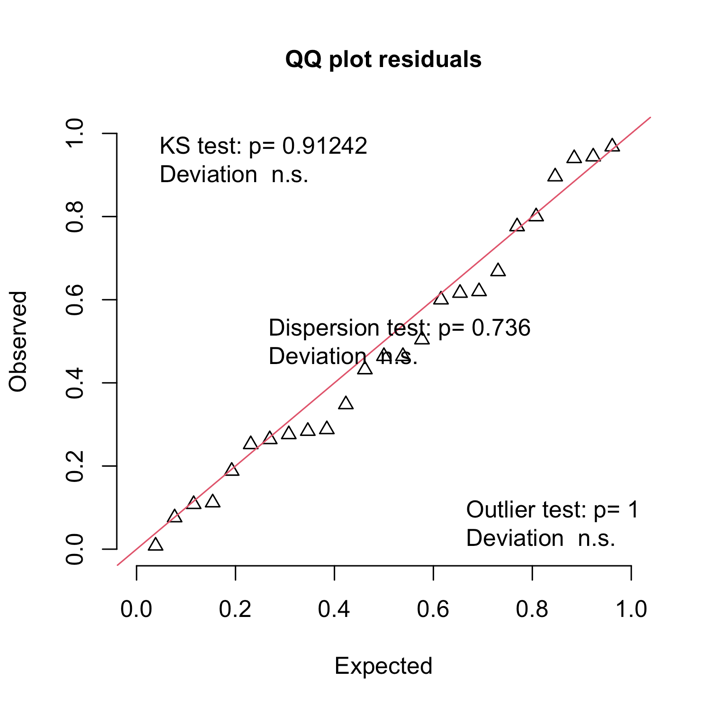

**A**

**B**

**Figure S2.** Residual plots for A) GLM predicting total Chao2 richness based on parasite genera detected per island population and B) GLM predicting total Chao2 richness based on parasite genera detected per island population per transmission mode.

**Table S4.** Model selection results for GLM predicting total Chao2 richness based on parasite genera detected per island population. Each row represents an individual model and includes parameter estimates (intercept and coefficients), degrees of freedom (df), log-likelihood (logLik), Akaike Information Criterion corrected for small sample size (AICc), delta AICc (ΔAICc), and Akaike weights. NA values indicate that the parameter was not included in the model. Area = log_10_ Island Area (km²), Isolation = log_10_ isolation from the Australian continent (km).

| Intercept | Area | Latitude | Isolation | | Df | Log Likelihood | | AICc | ΔAICc | Weight |
| --- | --- | --- | --- | --- | --- | --- | --- | --- | --- | --- |
| 5.64 | NA | 0.01 | | -0.38 | 4 | -956.72 | 1923.45 | | 0.00 | 0.51 |
| 5.58 | -0.01 | 0.01 | | -0.36 | 5 | -955.18 | 1923.51 | | 0.06 | 0.49 |
| 5.01 | -0.03 | NA | | -0.31 | 4 | -971.30 | 1952.61 | | 29.16 | 0.00 |
| 5.04 | NA | NA | | -0.35 | 3 | -978.82 | 1964.79 | | 41.34 | 0.00 |
| 2.92 | -0.05 | NA | | NA | 3 | -1021.79 | 2050.72 | | 127.27 | 0.00 |
| 2.96 | -0.05 | 0.00 | | NA | 4 | -1021.41 | 2052.83 | | 129.38 | 0.00 |
| 2.75 | NA | 0.01 | | NA | 3 | -1040.68 | 2088.50 | | 165.05 | 0.00 |
| 2.56 | NA | NA | | NA | 2 | -1044.51 | 2093.57 | | 170.12 | 0.00 |
| 5.64 | NA | 0.01 | | -0.38 | 4 | -956.72 | 1923.45 | | 0.00 | 0.51 |
| 5.58 | -0.01 | 0.01 | | -0.36 | 5 | -955.18 | 1923.51 | | 0.06 | 0.49 |
| 5.01 | -0.03 | NA | | -0.31 | 4 | -971.30 | 1952.61 | | 29.16 | 0.00 |
| 5.04 | NA | NA | | -0.35 | 3 | -978.82 | 1964.79 | | 41.34 | 0.00 |
| 2.92 | -0.05 | NA | | NA | 3 | -1021.79 | 2050.72 | | 127.27 | 0.00 |
| 2.96 | -0.05 | 0.00 | | NA | 4 | -1021.41 | 2052.83 | | 129.38 | 0.00 |
| 2.75 | NA | 0.01 | | NA | 3 | -1040.68 | 2088.50 | | 165.05 | 0.00 |
| 2.56 | NA | NA | | NA | 2 | -1044.51 | 2093.57 | | 170.12 | 0.00 |

**Table S5.** Model selection results for GLM predicting total Chao2 richness based on parasite genera detected per island population. Each row represents an individual model and includes parameter estimates (intercept and coefficients), degrees of freedom (df), log-likelihood (logLik), Akaike Information Criterion corrected for small sample size (AICc), delta AICc (ΔAICc), and Akaike weights. NA values indicate that the parameter was not included in the model. Area = log_10_ Island Area (km²), Isolation = log_10_ isolation from the Australian continent (km), TS = Transmission Strategy.

| Intercept | | Area | Latitude | Isolation | TS | Area x TS | Isolation x TS | Df | Log Likelihood | AICc | ΔAICc | Weight |
| --- | --- | --- | --- | --- | --- | --- | --- | --- | --- | --- | --- | --- |
| 1.83 | 0.01 | | 0.00 | -0.12 | + | + | + | 11 | -1761.49 | 3549.45 | 0.00 | 0.90 |
| 1.60 | 0.01 | | NA | -0.09 | + | + | + | 10 | -1765.06 | 3553.79 | 4.34 | 0.10 |
| 1.64 | NA | | NA | -0.09 | + | NA | + | 7 | -1788.22 | 3592.21 | 42.76 | 0.00 |
| 1.75 | NA | | 0.00 | -0.10 | + | NA | + | 8 | -1787.46 | 3593.25 | 43.79 | 0.00 |
| 1.62 | 0.00 | | NA | -0.09 | + | NA | + | 8 | -1788.09 | 3594.51 | 45.06 | 0.00 |
| 1.76 | 0.01 | | 0.00 | -0.11 | + | NA | + | 9 | -1786.88 | 3594.72 | 45.26 | 0.00 |
| 4.06 | 0.01 | | 0.00 | -0.42 | + | + | NA | 9 | -1816.44 | 3653.84 | 104.38 | 0.00 |
| 3.87 | 0.01 | | NA | -0.41 | + | + | NA | 8 | -1818.98 | 3656.28 | 106.83 | 0.00 |
| 4.18 | NA | | 0.00 | -0.43 | + | NA | NA | 6 | -1832.86 | 3679.03 | 129.58 | 0.00 |
| 4.06 | NA | | NA | -0.42 | + | NA | NA | 5 | -1834.42 | 3679.77 | 130.32 | 0.00 |
| 4.06 | -0.01 | | NA | -0.42 | + | NA | NA | 6 | -1834.02 | 3681.35 | 131.89 | 0.00 |
| 4.17 | 0.00 | | 0.00 | -0.43 | + | NA | NA | 7 | -1832.84 | 3681.45 | 132.00 | 0.00 |
| 4.53 | NA | | 0.01 | -0.44 | NA | NA | NA | 4 | -1935.32 | 3879.25 | 329.80 | 0.00 |
| 4.48 | -0.01 | | 0.00 | -0.43 | NA | NA | NA | 5 | -1934.66 | 3880.24 | 330.79 | 0.00 |
| 4.29 | -0.01 | | NA | -0.41 | NA | NA | NA | 4 | -1937.01 | 3882.63 | 333.18 | 0.00 |
| 4.31 | NA | | NA | -0.42 | NA | NA | NA | 3 | -1939.37 | 3885.09 | 335.64 | 0.00 |
| 0.77 | 0.00 | | -0.01 | NA | + | + | NA | 8 | -1952.19 | 3922.70 | 373.25 | 0.00 |
| 0.93 | 0.01 | | NA | NA | + | + | NA | 7 | -1961.91 | 3939.60 | 390.15 | 0.00 |
| 1.06 | -0.05 | | -0.01 | NA | + | NA | NA | 6 | -1972.41 | 3958.13 | 408.67 | 0.00 |
| 1.23 | -0.04 | | NA | NA | + | NA | NA | 5 | -1985.14 | 3981.21 | 431.76 | 0.00 |
| 0.86 | NA | | -0.01 | NA | + | NA | NA | 5 | -2001.58 | 4014.09 | 464.64 | 0.00 |
| 0.97 | NA | | NA | NA | + | NA | NA | 4 | -2004.32 | 4017.25 | 467.80 | 0.00 |
| 1.42 | -0.06 | | -0.01 | NA | NA | NA | NA | 4 | -2054.09 | 4116.79 | 567.34 | 0.00 |
| 1.60 | -0.05 | | NA | NA | NA | NA | NA | 3 | -2063.68 | 4133.71 | 584.26 | 0.00 |
| 1.21 | NA | | 0.00 | NA | NA | NA | NA | 3 | -2089.52 | 4185.39 | 635.94 | 0.00 |
| 1.30 | NA | | NA | NA | NA | NA | NA | 2 | -2090.63 | 4185.43 | 635.98 | 0.00 |
| 1.83 | 0.01 | | 0.00 | -0.12 | + | + | + | 11 | -1761.49 | 3549.45 | 0.00 | 0.90 |
| 1.60 | 0.01 | | NA | -0.09 | + | + | + | 10 | -1765.06 | 3553.79 | 4.34 | 0.10 |
| 1.64 | NA | | NA | -0.09 | + | NA | + | 7 | -1788.22 | 3592.21 | 42.76 | 0.00 |
| 1.75 | NA | | 0.00 | -0.10 | + | NA | + | 8 | -1787.46 | 3593.25 | 43.79 | 0.00 |
| 1.62 | 0.00 | | NA | -0.09 | + | NA | + | 8 | -1788.09 | 3594.51 | 45.06 | 0.00 |
| 1.76 | 0.01 | | 0.00 | -0.11 | + | NA | + | 9 | -1786.88 | 3594.72 | 45.26 | 0.00 |
| 4.06 | 0.01 | | 0.00 | -0.42 | + | + | NA | 9 | -1816.44 | 3653.84 | 104.38 | 0.00 |
| 3.87 | 0.01 | | NA | -0.41 | + | + | NA | 8 | -1818.98 | 3656.28 | 106.83 | 0.00 |
| 4.18 | NA | | 0.00 | -0.43 | + | NA | NA | 6 | -1832.86 | 3679.03 | 129.58 | 0.00 |
| 4.06 | NA | | NA | -0.42 | + | NA | NA | 5 | -1834.42 | 3679.77 | 130.32 | 0.00 |
| 4.06 | -0.01 | | NA | -0.42 | + | NA | NA | 6 | -1834.02 | 3681.35 | 131.89 | 0.00 |
| 4.17 | 0.00 | | 0.00 | -0.43 | + | NA | NA | 7 | -1832.84 | 3681.45 | 132.00 | 0.00 |
| 4.53 | NA | | 0.01 | -0.44 | NA | NA | NA | 4 | -1935.32 | 3879.25 | 329.80 | 0.00 |
| 4.48 | -0.01 | | 0.00 | -0.43 | NA | NA | NA | 5 | -1934.66 | 3880.24 | 330.79 | 0.00 |
| 4.29 | -0.01 | | NA | -0.41 | NA | NA | NA | 4 | -1937.01 | 3882.63 | 333.18 | 0.00 |
| 4.31 | NA | | NA | -0.42 | NA | NA | NA | 3 | -1939.37 | 3885.09 | 335.64 | 0.00 |
| 0.77 | 0.00 | | -0.01 | NA | + | + | NA | 8 | -1952.19 | 3922.70 | 373.25 | 0.00 |
| 0.93 | 0.01 | | NA | NA | + | + | NA | 7 | -1961.91 | 3939.60 | 390.15 | 0.00 |
| 1.06 | -0.05 | | -0.01 | NA | + | NA | NA | 6 | -1972.41 | 3958.13 | 408.67 | 0.00 |
| 1.23 | -0.04 | | NA | NA | + | NA | NA | 5 | -1985.14 | 3981.21 | 431.76 | 0.00 |
| 0.86 | NA | | -0.01 | NA | + | NA | NA | 5 | -2001.58 | 4014.09 | 464.64 | 0.00 |
| 0.97 | NA | | NA | NA | + | NA | NA | 4 | -2004.32 | 4017.25 | 467.80 | 0.00 |
| 1.42 | -0.06 | | -0.01 | NA | NA | NA | NA | 4 | -2054.09 | 4116.79 | 567.34 | 0.00 |
| 1.60 | -0.05 | | NA | NA | NA | NA | NA | 3 | -2063.68 | 4133.71 | 584.26 | 0.00 |
| 1.21 | NA | | 0.00 | NA | NA | NA | NA | 3 | -2089.52 | 4185.39 | 635.94 | 0.00 |
| 1.30 | NA | | NA | NA | NA | NA | NA | 2 | -2090.63 | 4185.43 | 635.98 | 0.00 |
| 1.83 | 0.01 | | 0.00 | -0.12 | + | + | + | 11 | -1761.49 | 3549.45 | 0.00 | 0.90 |
| 1.60 | 0.01 | | NA | -0.09 | + | + | + | 10 | -1765.06 | 3553.79 | 4.34 | 0.10 |
| 1.64 | NA | | NA | -0.09 | + | NA | + | 7 | -1788.22 | 3592.21 | 42.76 | 0.00 |
| 1.75 | NA | | 0.00 | -0.10 | + | NA | + | 8 | -1787.46 | 3593.25 | 43.79 | 0.00 |
| 1.62 | 0.00 | | NA | -0.09 | + | NA | + | 8 | -1788.09 | 3594.51 | 45.06 | 0.00 |
| 1.76 | 0.01 | | 0.00 | -0.11 | + | NA | + | 9 | -1786.88 | 3594.72 | 45.26 | 0.00 |
| 4.06 | 0.01 | | 0.00 | -0.42 | + | + | NA | 9 | -1816.44 | 3653.84 | 104.38 | 0.00 |
| 3.87 | 0.01 | | NA | -0.41 | + | + | NA | 8 | -1818.98 | 3656.28 | 106.83 | 0.00 |
| 4.18 | NA | | 0.00 | -0.43 | + | NA | NA | 6 | -1832.86 | 3679.03 | 129.58 | 0.00 |
| 4.06 | NA | | NA | -0.42 | + | NA | NA | 5 | -1834.42 | 3679.77 | 130.32 | 0.00 |
| 4.06 | -0.01 | | NA | -0.42 | + | NA | NA | 6 | -1834.02 | 3681.35 | 131.89 | 0.00 |
| 4.17 | 0.00 | | 0.00 | -0.43 | + | NA | NA | 7 | -1832.84 | 3681.45 | 132.00 | 0.00 |
| 4.53 | NA | | 0.01 | -0.44 | NA | NA | NA | 4 | -1935.32 | 3879.25 | 329.80 | 0.00 |
| 4.48 | -0.01 | | 0.00 | -0.43 | NA | NA | NA | 5 | -1934.66 | 3880.24 | 330.79 | 0.00 |
| 4.29 | -0.01 | | NA | -0.41 | NA | NA | NA | 4 | -1937.01 | 3882.63 | 333.18 | 0.00 |
| 4.31 | NA | | NA | -0.42 | NA | NA | NA | 3 | -1939.37 | 3885.09 | 335.64 | 0.00 |
| 0.77 | 0.00 | | -0.01 | NA | + | + | NA | 8 | -1952.19 | 3922.70 | 373.25 | 0.00 |
| 0.93 | 0.01 | | NA | NA | + | + | NA | 7 | -1961.91 | 3939.60 | 390.15 | 0.00 |
